# Dynamically actuated reconfigurable topographical surface enables active control of implant-associated infections

**DOI:** 10.64898/2026.06.29.735318

**Authors:** Mohammad Asadi Tokmedash, Jihun Lee, J. Scott VanEpp, Sungmin Nam, Jouha Min

## Abstract

Implant-associated infections are driven by bacterial biofilm formation and remain difficult to eradicate using conventional antibiotic-based strategies. Here, we present a dynamically actuated reconfigurable topographical surface (DARTS) that integrates intrinsically bactericidal nanoscale surface topography with programmable mechanical actuation to achieve durable, antibiotic-free infection control. Using a scalable bottom-up nanofabrication strategy, we generate tunable wrinkled MXene topographies that exhibit contact-mediated bactericidal activity against both Gram-positive and Gram-negative bacteria without chemical leaching. Integration with a soft robotic actuator enables reversible modulation of surface geometry, which synergistically enhances bacterial removal and killing, resulting in near-complete disruption of mature biofilms. Dynamic actuation further sensitizes released bacteria to antibiotic treatment. In a mouse subcutaneous implant infection model, DARTS with actuation achieves sustained suppression of bacterial burden and markedly improves host tissue outcomes. Remote, noninvasive actuation using near-infrared laser stimulation further highlights the translational potential of this platform for implantable antibacterial applications.

**Significance Statement:** Implant infections are difficult to treat because bacteria form biofilms that protect them from antibiotics and the immune system. Current materials often rely on chemical release, which can lose effectiveness over time. Here, we present a new surface that both kills bacteria and removes them. The surface uses nanoscale features to physically damage bacterial cells, while dynamic motion clears attached bacteria and biofilms. This allows continuous, chemical-free control of infection. In a mouse implant model, the system greatly reduced bacterial burden and improved tissue healing. This work introduces a new way to control bacteria using dynamic surface design and could be applied not only to medical implants but also to environmental, food, textile, and marine systems.

## INTRODUCTION

Implant-associated infection remains one of the most frequent and severe complications in modern medicine,^1^ accounting for over 25% of all healthcare-associated infections (HAIs) in the United States.^2^ These infections frequently result in implant failure, with reported rates exceeding 25% for artificial heart valves,^3^ up to 35% for post-mastectomy breast implants,^4^ and 74% for revision hip arthroplasties.^5^ Clinically, implant infections often necessitate multiple revision surgeries and can progress to chronic inflammation, sepsis, or death.^6^ The economic burden is equally substantial; HAIs affect approximately 440,000 adult inpatients annually and generate nearly $10 billion in health care costs across major infection categories, including surgical site infections, ventilator-associated pneumonia, catheter-associated urinary tract infections, and central line-associated bloodstream infections.^7^ Despite these consequences, infection risk has historically been underemphasized in the design of implantable materials and devices, and remains insufficiently addressed in emerging tissue engineering strategies.^8^

Most implant-associated infections are initiated by bacterial adhesion and subsequent biofilm formation.^9–11^ Biofilms are highly organized multicellular communities that exhibit extreme tolerance to antibiotics and host immune defenses, rendering established infections difficult to eradicate.^12–14^ Current prevention strategies primarily rely on antimicrobial surface coatings or local drug systems that release antibiotics^15–17^ or metal-based agents such as silver, zinc oxide, copper, or gold nanoparticles.^18–22^ While these approaches can reduce early-stage bacterial colonization, they suffer from fundamental limitations, including finite efficacy due to agent depletion, narrow therapeutic windows, and concerns regarding cytotoxicity and systemic exposure.^23–28^ Moreover, the rapid rise of antibiotic-resistant pathogens, including methicillin-resistant *Staphylococcus aureus* (MRSA) and extended-spectrum β-lactamase-producing bacteria, has underscored the urgency of developing agent-free strategies that do not exert selective pressure for resistance.

Surface topographical modification has emerged as a promising non-chemical strategy for regulating bacterial attachment and biofilm formation. By engineering surface features at the nano- and micro-scale, bacterial adhesion can be suppressed and, in some cases, bacterial membranes can be mechanically disrupted, enabling antifouling or bactericidal effects without relying on the release of antimicrobial agents.^29–35^ Nanoscale topographies inspired by natural antibacterial surfaces, such as cicada and dragonfly wings, have been shown to induce physical damage to bacterial envelopes through direct mechanical interactions.^36–41^ At larger length scales, microscale surface features can also reduce bacterial attachment by modulating cell–material interactions and limiting stable bacterial colonization.^42–44^

However, most antibacterial topographies are inherently static, and their effectiveness often diminishes over clinically relevant timescales. As bacteria adapt, proliferate, and form mature biofilms, static surface features are eventually overcome, resulting in reduced long-term antibacterial performance.^45, 46^ To address this limitation, dynamic surface topographies have been explored using mechanical stretching,^47, 48^ pneumatic actuation,^49, 50^ electrical stimulation,^47^ or magnetic fields.^51^ While these approaches can disrupt early-stage bacterial attachment and reduce fouling, they generally lack intrinsic bactericidal activity and frequently require adjunctive antibiotic treatment. Moreover, many dynamic topography platforms rely on conventional lithographic fabrication, limiting them to small scales and restricting translation to implant-relevant geometries. In parallel, advances in soft robotics have enabled programmable mechanical actuation in vivo^52^, yet these technologies have not been applied to the dynamic reconfiguration of surface topography at the cell-material interface. Collectively, these limitations underscore a critical unmet need for scalable, implant-compatible platforms that integrate intrinsically bactericidal surface topographies with programmable, dynamic actuation to achieve durable, antibiotic-free antibacterial performance in vivo.

Here, we introduce a novel antibacterial platform based on dynamically actuated reconfigurable topographical surfaces (DARTS) (**Fig. 1**). This system integrates bottom-up nanofabrication with soft robotic actuation to enable programmable and reversible modulation of surface geometry. By leveraging purely physical mechanisms, DARTS provides a chemical-free strategy to regulate bacteria–surface interactions on soft, implant-relevant materials. We demonstrate robust antibacterial performance across a panel of clinically relevant Gram-positive and Gram-negative bacterial species, with consistent efficacy validated in both in vitro and in vivo settings.

**Fig. 1.**
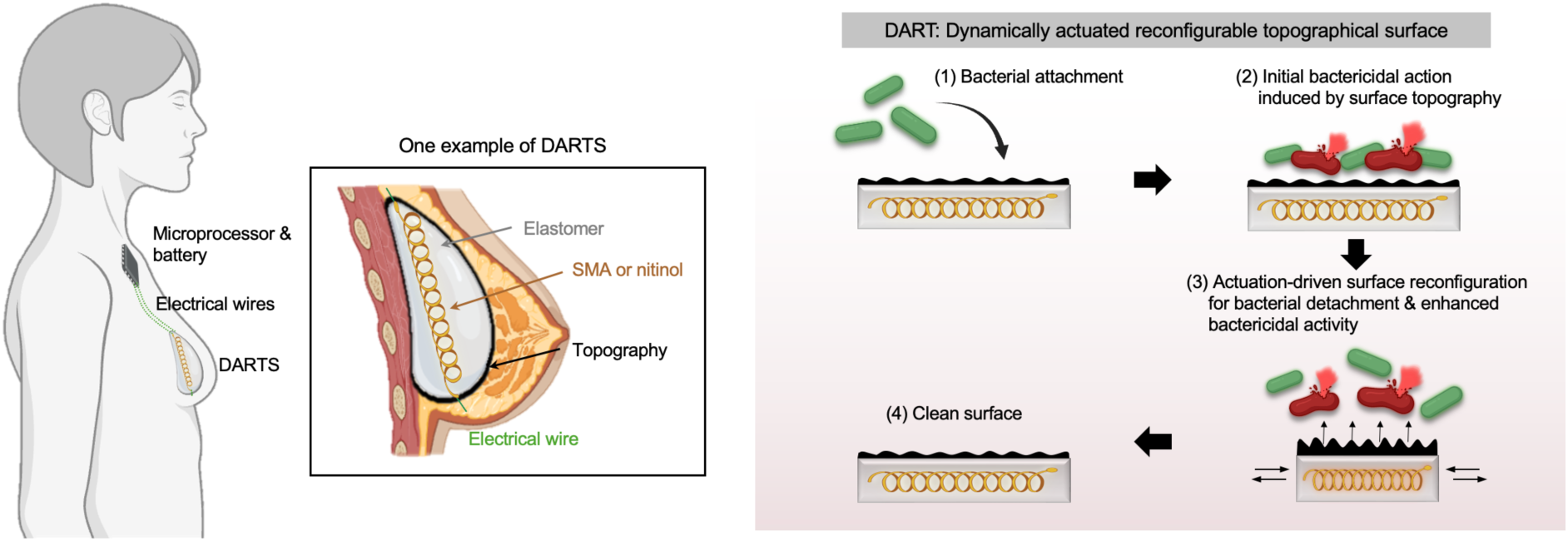
Concept and mechanism of DARTS. DARTS integrates bactericidal surface topography with programmable actuation to disrupt bacterial attachment and promote biofilm removal, enabling durable, chemical-free antibacterial performance at implant interfaces.

## RESULTS

### Working principle of dynamic surface topography

The antibacterial function of DARTS is governed by two distinct yet complementary mechanisms: static surface topography and dynamic mechanical actuation (Fig. 1). Under static conditions, nano- and micro-scale surface features regulate bacteria–surface interactions by reducing effective contact area and inhibiting the formation of stable adhesions. Densely packed, high-curvature topographies suppress bacterial attachment and generate localized mechanical stresses that compromise bacterial membranes, resulting in bactericidal activity and reduced surface colonization. Dynamic mechanical actuation further modulates these interactions by introducing controlled deformation at the bacteria–material interface. This deformation promotes the detachment of weakly or transiently adhered bacteria while simultaneously amplifying mechanically induced membrane damage in bacteria that remain attached. In the following sections, we first examine the antibacterial effects of surface topography alone under static conditions. We then investigate how the integration of surface topography with dynamic actuation synergistically enhances both bacterial removal and bactericidal efficacy.

### Bottom-up nanofabrication of tunable wrinkled topographies

To engineer surface topographies spanning the nano- to micro-scale, we employed our previously developed bottom-up nanofabrication strategy (**Fig. 2A**).^53^ Alternating layers of two-dimensional Ti_3_C_2_T_x_ MXene nanosheets and a cationic counterpart were deposited onto biaxially pre-strained polystyrene (PS) substrates via electrostatic layer-by-layer (LbL) assembly, yielding conformal, mechanically stiff multilayer coatings. Consistent with prior reports,^53–56^ the total coating thickness increased linearly with the number of deposited bilayers (**Supplementary Fig. 1B-D**). Wrinkled surface topographies were subsequently generated by thermally induced biaxial contraction of the PS substrate. Substrate contraction imposed in-plane compressive stresses on the stiff multilayer coating, leading to elastic buckling and the formation of isotropic wrinkled architectures. This process produced well-defined surface features spanning multiple length scales. Scanning electron microscopy (SEM) revealed highly ordered wrinkle patterns with sharp, high-curvature features ranging from the nanoscale to the micron scale (**Fig. 2B**). Quantitative analysis revealed that both wrinkle wavelength and amplitude scaled linearly with bilayer number (**Fig. 2C**; **Supplementary Fig. 1E**). Specifically, the wrinkle wavelength increased from approximately 230 to 2700 nm, while the amplitude increased from approximately 180 to 1250 nm as the coating thickness increased from 1 to 20 bilayers (**Fig. 2C**). These scaling relationships are consistent with classical buckling theory^57–59^ (**Supplementary Note**) and confirm precise, predictable, and scalable control over surface geometry using this fabrication framework.

**Fig. 2.**
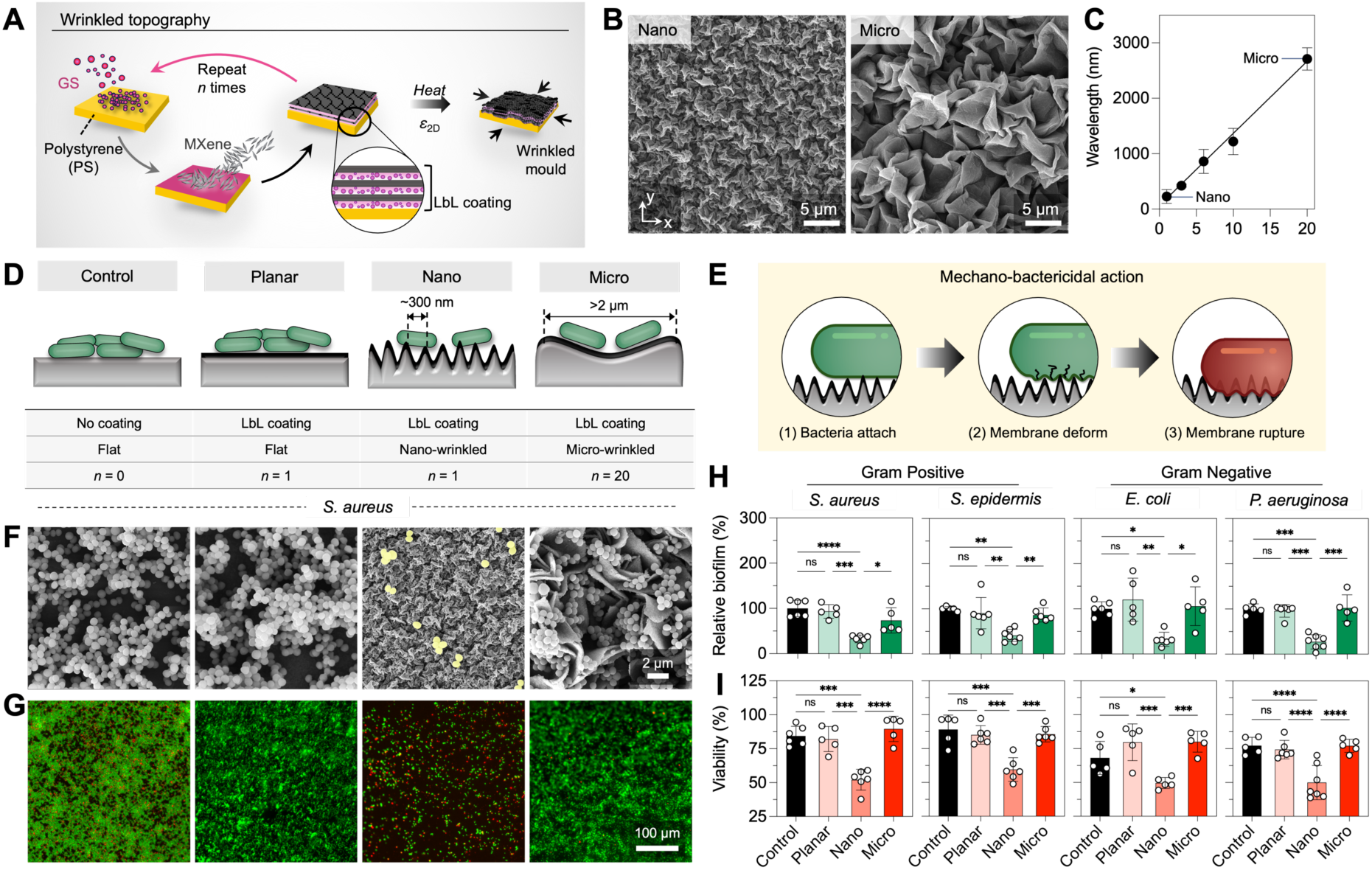
Fabrication, characterization, and antibacterial performance of wrinkled MXene topographies. **(A)** Schematic of the fabrication of wrinkled MXene multilayers on PS films. **(B)** Representative SEM images of the resulting surface topographies. **(C)** Wrinkle wavelength as a function of MXene bilayer number (n ≥ 3). **(D)** Schematic illustrating preferential bacterial attachment to microscale grooves and nanoscale features. **(E)** Schematic of nanoscale topography–induced bactericidal interactions with bacterial membranes. **(F)** SEM and **(G)** fluorescence images of *S. aureus* biofilms on Control, Planar, Nano, and Micro surfaces. Quantification of **(H)** relative biofilm coverage and **(I)** bacterial viability across multiple bacterial species (n = 5–6). Statistical analysis was performed using ordinary two-way ANOVA with Tukey’s post hoc multiple-comparison test; ns (not significant), *P < 0.05, **P < 0.01, ***P < 0.001, and ****P < 0.0001. Data are presented as mean ± S.D.

### In vitro evaluation for topography-driven bactericidal activity

We then evaluated how surface topography influences antibacterial activity. Four conditions were examined: Control (uncoated flat PS), Planar (flat MXene-coated PS, 1 bilayer), Nano (nano-wrinkled MXene-coated PS, 1 bilayer; λ < 400 nm), and Micro (micro-wrinkled MXene-coated PS, 20 bilayers; λ > 2 µm), covering a wide range of topographical length scales relevant to bacterial attachment and survival (**Fig. 2D** and **E**). Comparisons between the Control and Planar surfaces enabled assessment of antibacterial effects attributable to surface chemistry, whereas comparisons among the Planar, Nano, and Micro surfaces isolated the role of surface topography independent of chemistry. The engineered surfaces were challenged with multiple bacterial species, including the Gram-positive *Staphylococcus aureus* (*S. aureus*) and *Staphylococcus epidermidis* (*S. epidermidis*), as well as Gram-negative *Escherichia coli* (*E. coli*) and *Pseudomonas aeruginosa* (*P. aeruginosa*). SEM imaging revealed pronounced differences in bacterial adhesion and morphology across the four conditions (**Fig. 2F**; **Supplementary Fig. 2A**). Both Control and Planar surfaces supported dense bacterial colonization and biofilm formation, with bacteria maintaining intact morphology indicative of healthy growth. In contrast, the Nano surface exhibited markedly reduced bacterial attachment. By comparison, the Micro surface promoted bacterial accumulation, particularly within recessed micron-scale regions that appeared to accommodate bacterial attachment. Live/dead fluorescence imaging further supported these observations. Control, Planar, and Micro surfaces were densely populated with SYTO9-positive (live, green) bacteria, whereas the Nano surface exhibited sparse bacterial coverage dominated by propidium iodide-positive (dead, red) cells (**Fig. 2G**; **Supplementary Fig. 2B**). These qualitative trends were reflected in quantitative analyses of biofilm formation and bacterial viability. Across all tested species, biofilm formation decreased with decreasing topographical feature size, with the Nano surface consistently exhibiting the lowest biofilm coverage **(Fig. 2H)**. Likewise, viability measurements showed that the Nano surface significantly suppressed bacterial survival across both Gram-positive and Gram-negative species, whereas Control, Planar, and Micro surfaces largely preserved bacterial viability **(Fig. 2I)**.

To rule out the possibility that the observed antibacterial activity originated from chemical leaching rather than physical surface architecture, we conducted a non-contact antibacterial assay (**Supplementary Fig. 3**). Bacteria were cultured in the presence of the Nano and Micro surfaces without direct contact. A standard gentamicin sulfate antibiotic disk was included as a positive control and produced a clear zone of growth inhibition in the surrounding area. In contrast, neither the Nano nor the Micro surfaces generated detectable inhibition zones, even in close proximity to the coated surfaces. These results suggest that bacterial killing by the MXene-coated surfaces arises from contact-mediated, topography-driven mechanisms rather than from leached antimicrobial agents.

### Integration with a soft actuator for dynamic topographical modulation

After establishing the antibacterial effects of surface topography, we next integrated it with dynamic actuation. To achieve this, we employed a soft robotic actuator designed to generate controlled and repeatable contractile deformation (**Fig. 3A**). The actuator consists of a shape memory alloy (SMA) coil embedded within a soft elastomer matrix (Ecoflex). During fabrication, the SMA coil was mechanically stretched to an elongated state and subsequently encapsulated within the Ecoflex matrix. Upon electrical stimulation, Joule heating induces a martensite-to-austenite phase transformation in the SMA, driving recovery toward its original coiled geometry (**Fig. 3B**). This shape recovery generates a contractile force that compresses the surrounding elastomer and the overlying wrinkled surface layer, thereby dynamically modulating surface topography. Upon removal of the electrical input, the SMA cools and the elastic restoring force of the elastomer returns the actuator to its initial state, enabling reversible and repeatable topographical reconfiguration.

**Fig. 3.**
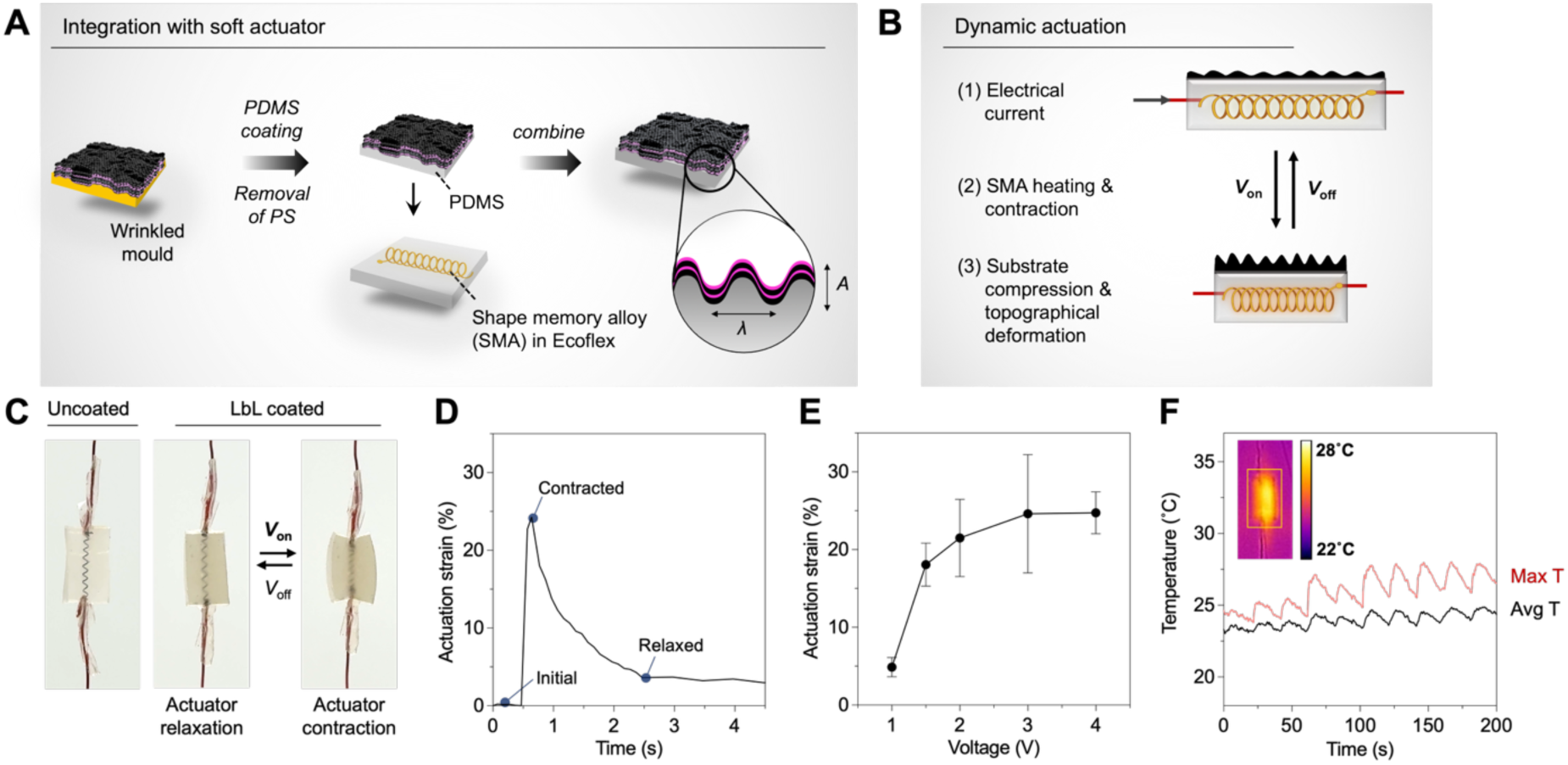
Fabrication, characterization, and actuation of DARTS. **(A)** Schematic of DARTS fabrication via transfer printing of wrinkled MXene topographies onto an SMA-based soft actuator through an intermediate PDMS layer. **(B)** Schematic of dynamic actuation driven by electrically induced SMA heating and contraction. **(C)** Optical images of DARTS before and during actuation, showing reversible deformation. **(D)** Cyclic actuation behavior of DARTS. **(E)** Actuation strain as a function of applied voltage (n = 5). **(F)** Infrared thermal imaging and corresponding temperature profiles during actuation. Data are presented as mean ± S.D.

To integrate the wrinkled MXene multilayer coatings with the soft actuator, we initially attempted direct deposition of the multilayers onto the actuator surface. However, the low elastic modulus of the actuator was insufficient to induce elastic buckling of the stiff coating, resulting in shallow and poorly defined surface features with minimal nanoscale wrinkling (**Supplementary Fig. 4**). To overcome this limitation, wrinkled MXene multilayers were first fabricated on biaxially pre-strained PS substrates, as described above, and subsequently transfer-printed onto a thin PDMS layer (∼400 µm thick), which served as an intermediate elastomeric support. This transfer-printing strategy successfully preserved nano-, and micron-scale features with high fidelity (**Supplementary Fig. 5**). Among the surface topographies, the Nano surface was selected for its superior intrinsic antibacterial performance and integrated into the Ecoflex-embedded SMA actuator to form DARTS.

We then evaluated the actuation performance of DARTS. Similar to the uncoated actuator and the PDMS-coated actuator, DARTS exhibited robust contraction upon voltage application and reliably returned to its initial state upon removal of the electrical input (**Fig. 3C**; **Supplementary Fig. 6**).

These results indicate that integration of the wrinkled surface layer does not compromise actuator performance and confirm highly repeatable actuation behavior. The magnitude of contraction was precisely tuned by the applied voltage, producing actuation strains of up to ∼25% (**Fig. 3D** and **E**). Thermal imaging during actuation revealed localized temperature increases confined to the central region of the actuator corresponding to the SMA, while both the maximum and average temperatures across the elastomeric body remained within physiologically relevant ranges (**Fig. 3F**; **Supplementary Fig. 7**). Notably, these temperature profiles were consistently maintained within physiological limits even under repeated actuation cycles and were comparable to those observed for uncoated and PDMS-coated actuators.

### In vitro evaluation of the synergistic effects of topography and dynamic actuation

We next evaluated the combined effects of surface topography and dynamic actuation on bacterial adhesion and biofilm stability. Biofilms of *S. aureus, S. epidermidis, E. coli*, and *P. aeruginosa* were first allowed to form undisturbed on DARTS surfaces for 24-48 hours to establish mature biofilms, after which dynamic actuation was applied. To identify effective actuation conditions, we varied both the number of actuation cycles and the magnitude of applied strain from 0 to 20. The extent of biofilm disruption increased with both parameters, demonstrating a clear dependence on actuation intensity **(Fig. 4B; Supplementary Fig. 8)**. At the highest condition tested (20 cycles and 20% strain), dynamic actuation led to near-complete disruption and removal of adherent biofilms, leaving only sparse residual debris on the surface **(Fig. 4C)**. In contrast, flat control surfaces lacking engineered topography supported robust bacterial adhesion and dense biofilm formation, consistent with earlier observations **(Fig. 4A** and **B**; **Fig. 1G-I)**.

**Fig. 4.**
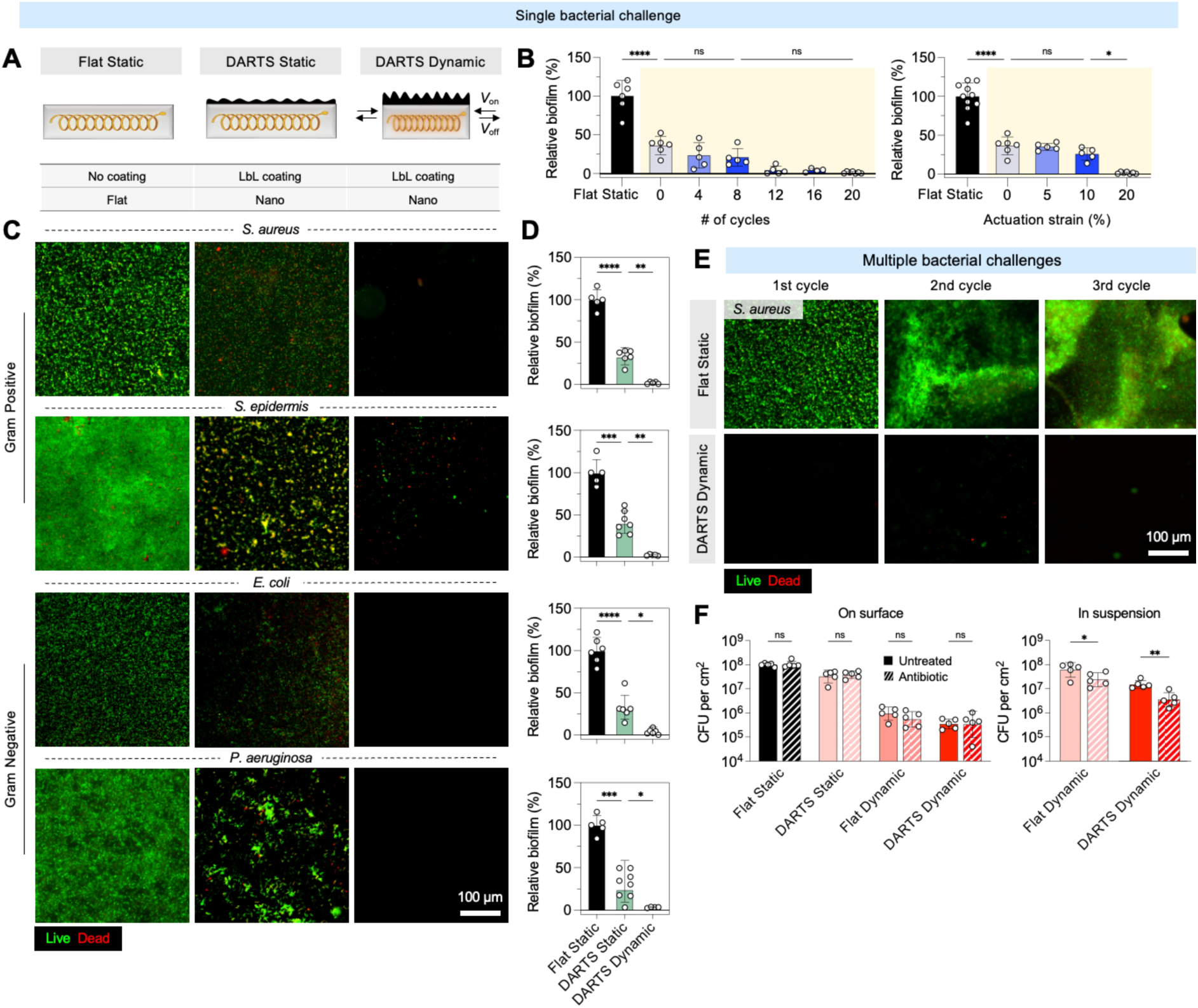
Synergistic antibiofilm activity of surface topography and dynamic actuation. **(A)** Schematic illustrating experimental design. **(B)** Dependence of biofilm disruption on actuation cycle number and strain magnitude (n = 5–10). **(C)** Representative live/dead fluorescence images of biofilms formed by multiple bacterial species under selected surface topography and actuation conditions. **(D)** Quantitative analysis of biofilm removal corresponding to the conditions shown in (C) (n = 5–8). **(E)** Biofilm removal performance under repeated bacterial contamination cycles. **(F)** CFU analysis of bacteria recovered from the surface and from the culture medium (in suspension) following dynamic actuation, with and without antibiotic treatment (n = 5). Statistical analysis was performed using ordinary two-way ANOVA with Tukey’s post hoc multiple-comparison test for (B) and (D), and two-sided nonparametric t-tests for (F); ns (not significant), *P < 0.05, **P < 0.01, ***P < 0.001, and ****P < 0.0001. Data are presented as mean ± S.D.

Under static conditions without actuation, DARTS exhibited reduced biofilm coverage relative to flat controls, reflecting its intrinsic topography-driven antibacterial activity **(Fig. 4C)**. However, a measurable population of viable bacteria remained adherent, indicating that static topographical cues alone were insufficient to achieve complete biofilm removal. We further assessed whether this performance was maintained under repeated contamination events (**Fig. 4E**). When subjected to three consecutive bacterial contamination cycles, flat control surfaces exhibited progressively denser and more stable *S. aureus* biofilms. In contrast, actuated DARTS consistently achieved near-complete biofilm removal across all cycles, highlighting its robustness against repeated bacterial challenge.

We further examined whether mechanical disruption of biofilms by DARTS enhances bacterial susceptibility to antibiotic treatment. Vancomycin hydrochloride, a clinically relevant glycopeptide antibiotic commonly used against Gram-positive biofilm-associated infections, was selected as a model antibiotic. S. aureus biofilms were first allowed to form on DARTS surfaces, after which the dynamic actuation was applied, and culture medium was replaced with vancomycin-containing medium. Bacterial viability was subsequently quantified by colony-forming unit (CFU) analysis.

Although dynamic actuation alone promoted biofilm dispersal on flat surfaces, the absence of bactericidal nanostructures resulted in substantially higher numbers of viable bacteria (Fig. 4F). In contrast, bacteria released from DARTS exhibited markedly reduced viability. These results indicate that DARTS not only facilitates efficient biofilm removal but also mechanically sensitizes bacteria upon release, thereby enhancing the effectiveness of antibiotic treatment.

### In vivo evaluation using a mouse subcutaneous implant model

To determine whether the antibacterial performance of DARTS observed in vitro translates in vivo, we employed a mouse subcutaneous implant infection model (**Fig. 5A**; **Supplementary Fig. 9**).

**Fig. 5.**
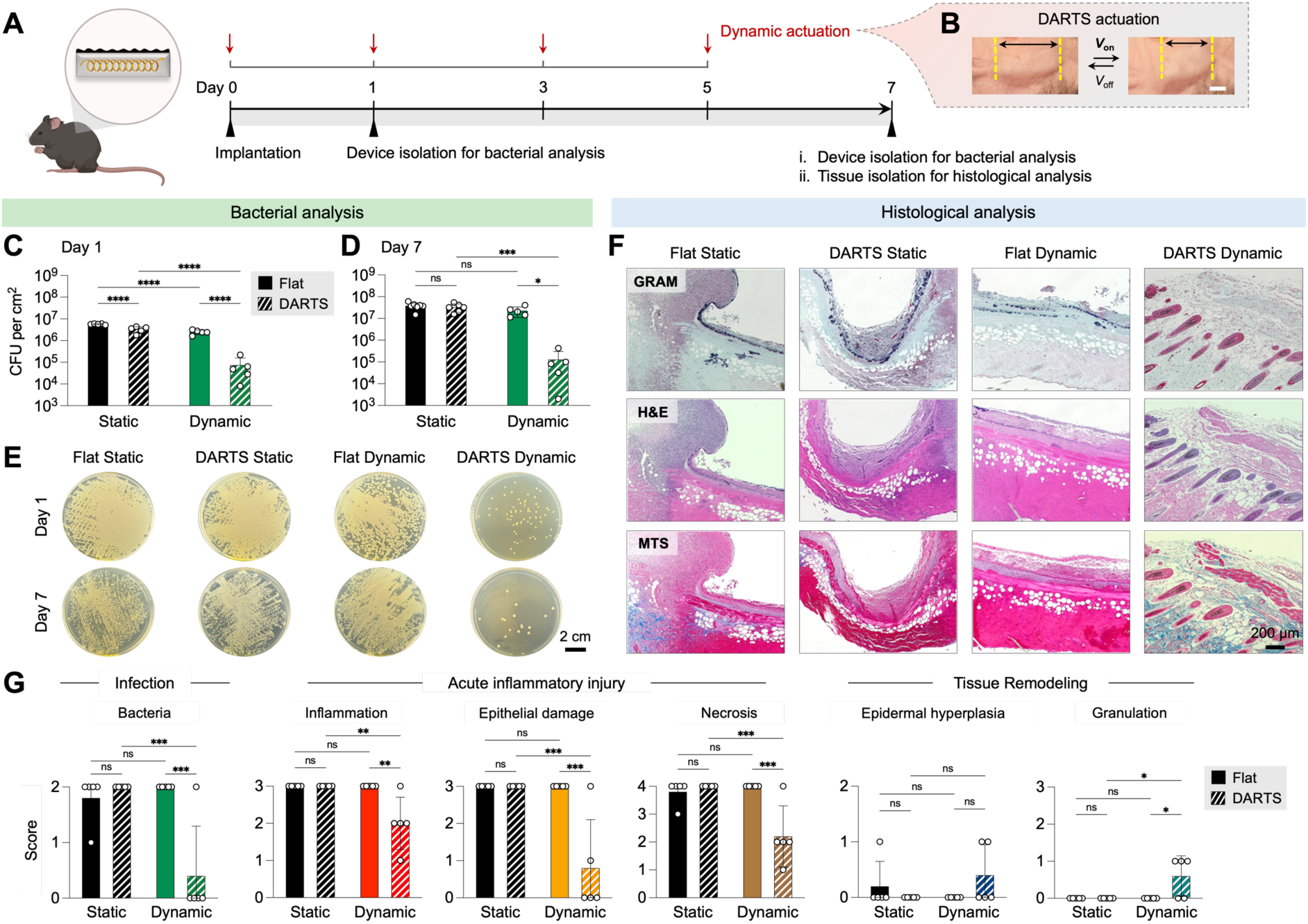
In vivo antibacterial performance of DARTS in a mouse subcutaneous implant infection model. **(A)** Schematic of the subcutaneous implant infection model and actuation protocol. **(B)** Photographs demonstrating in vivo DARTS actuation. **(C, D)** Quantification of bacterial burden recovered from implants at day 1 and day 7 post-implantation under different surface topography and actuation conditions (n = 5–7). **(E)** Representative agar plate images corresponding to the CFU quantification shown in (C, D). **(F)** Representative histological images of peri-implant tissues stained to assess bacterial colonization and host tissue response. **(G)** Blinded histological scoring of inflammation, tissue damage, necrosis, and bacterial burden (n = 5). Scale bar: 2mm in (B). Statistical analysis was performed using ordinary one-way ANOVA with Tukey’s post hoc multiple-comparison test; ns (not significant), *P < 0.05, **P < 0.01, ***P < 0.001, and ****P < 0.0001. Data are presented as mean ± S.D.

Early post-surgical contamination was simulated by implanting devices bearing pre-formed *S. aureus* biofilms, followed by dynamic actuation applied at multiple time points after implantation (**Fig. 5B**). Based on the actuation conditions that exhibited the strongest antibacterial performance in vitro (**Fig. 4**), a regimen of 20 actuation cycles at 20% actuation strain was selected for in vivo experiments. Implants were retrieved at early (day 1) and later (day 7) time points to assess bacterial burden and host tissue response. At day 1, implants incorporating either surface topography alone (DARTS Static) or dynamic actuation alone (Flat Dynamic) exhibited modest reductions in bacterial survival compared with flat, non-actuated controls (Flat Static), indicating partial antibacterial effects from each individual component (**Fig. 5C** and **E**). However, by day 7, bacterial burden increased in these groups, and differences relative to controls were largely diminished, demonstrating that neither surface topography nor dynamic actuation alone provided sustained antibacterial protection in vivo (**Fig. 5D** and **E**). In contrast, DARTS with actuation (DARTS Dynamic) consistently exhibited minimal bacterial survival at both day 1 and day 7 (**Fig. 5C-E**), highlighting that the synergistic antibacterial effect observed for DARTS in vitro is preserved in vivo.

To assess local infection severity and host tissue response, histological analyses were performed on tissue sections harvested 7 days after implantation (**Fig. 5F**). Gram staining revealed extensive bacterial colonization at the implant–tissue interface on flat surfaces, regardless of whether dynamic actuation was applied. Similarly, DARTS without actuation exhibited a substantial bacterial burden comparable to that of flat controls. In contrast, DARTS with actuation displayed a markedly reduced bacterial burden relative to all other conditions. Hematoxylin and eosin (H&E) and Masson’s trichrome staining (MTS) further revealed dense inflammatory cell infiltration and pronounced tissue damage surrounding the implant in the flat surface group. The DARTS Static and Flat Dynamic groups showed partial improvement; however, substantial inflammation and disrupted tissue architecture persisted, indicating incomplete infection control and a sustained host inflammatory response. In contrast, tissues adjacent to the DARTS Dynamic group exhibited well-preserved tissue architecture with minimal inflammatory infiltration. These qualitative observations were confirmed by blinded histological scoring (**Fig. 5G**), which showed significantly lower scores for inflammation, epithelial damage, necrosis, and bacterial burden in the DARTS Dynamic group, whereas the remaining groups exhibited comparably elevated scores across these parameters. Epidermal hyperplasia, indicative of chronic epithelial thickening, was minimal and comparable across all groups. Granulation tissue formation was generally low but modestly increased in the DARTS Dynamic group, likely due to early-stage tissue remodeling following effective suppression of infection.

### Remote-controllable actuation

Finally, as a proof of concept, we demonstrated remote actuation of DARTS using near-infrared (NIR) laser stimulation. NIR irradiation induced localized photothermal heating of the embedded SMA coil, resulting in controlled actuator contraction (**Fig. 6A**). Temperature measurements showed that the SMA reached approximately 38 °C during actuation, while the surrounding elastomer remained at lower temperatures within a physiologically relevant range (**Fig. 6B** and **C**). Importantly, this remote actuation was also successfully achieved in vivo (**Fig. 6D**). NIR laser irradiation effectively penetrated the skin and activated the implanted DARTS, producing controlled deformation without the need for direct electrical connections or wired interfaces. These results demonstrate that, beyond electrical stimulation, DARTS can be actuated using noninvasive, remote modalities such as NIR laser irradiation, highlighting its potential for translational, implantable antibacterial applications.

**Fig. 6.**
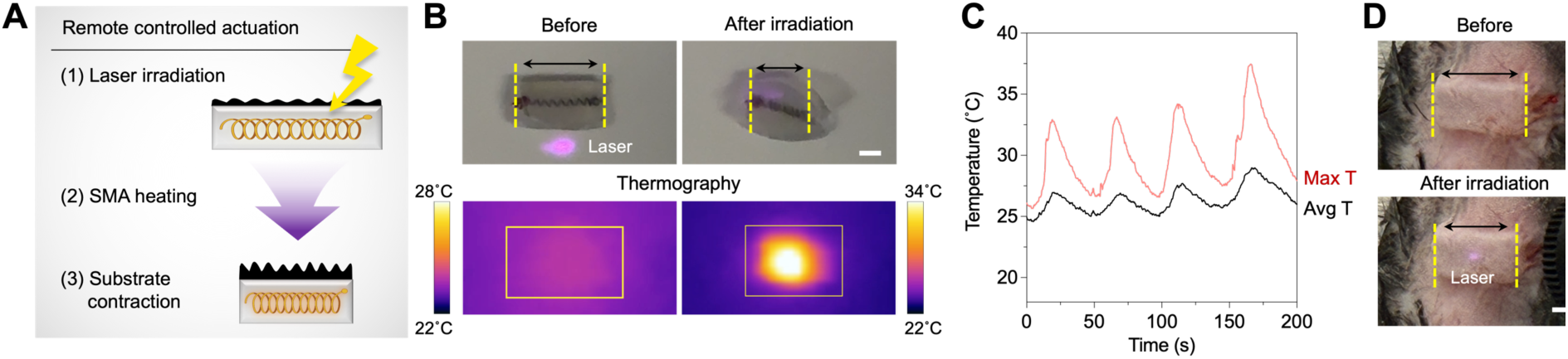
Remote actuation of DARTS via near-infrared laser stimulation. **(A**) Schematic of remote actuation in which near-infrared laser irradiation induces localized heating of the SMA, leading to actuator contraction and surface reconfiguration. **(B)** Optical and infrared thermal images of DARTS before and during laser irradiation. **(C)** Maximum and average temperature profiles during repeated laser actuation cycles. **(D)** Photographs demonstrating in vivo actuation of DARTS under remote stimulation. Scale bar, 2 mm in (B) and (D).

## DISCUSSION

This study introduces DARTS as a new materials strategy for infection-resistant biomaterials. In DARTS, nanoscale surface topography provides a geometry-driven antibacterial mechanism by suppressing bacterial adhesion and mechanically compromising cell membranes across both Gram-positive and Gram-negative species. However, as observed here and in prior studies^35, 60–63^, static topographical features alone are insufficient to sustain long-term antibacterial protection, as their efficacy diminishes over time due to progressive surface fouling by dead cells and extracellular matrix. To overcome this temporal limitation, DARTS incorporates dynamic actuation as a complementary mechanism, converting a static antibacterial surface into an actively renewable interface. Cyclic actuation generates transient shear and normal stresses that mechanically disrupt adherent bacteria and mature biofilms, leading to robust and repeatable biofilm removal across multiple bacterial species. Beyond physical clearing, this mechanical disruption compromises membrane integrity and disrupts biofilm structure, thereby increasing bacterial exposure to antibiotics. As a result, bacteria released from DARTS exhibit reduced viability and enhanced susceptibility to vancomycin compared to intact biofilms and bacteria detached from flat dynamic controls. These findings indicate that mechanical disruption induced by DARTS not only restores surface functionality but also synergistically enhances antibiotic efficacy in a manner that is difficult to achieve using static surfaces or chemical delivery alone.

This mechanically renewable antibacterial strategy translates effectively in vivo. In a mouse subcutaneous implant infection model, dynamically actuated DARTS implants consistently exhibited the lowest bacterial burden at both early and later time points, outperforming surfaces relying on topography or actuation alone. Histological analyses revealed preserved tissue architecture, reduced necrosis, and attenuated inflammatory infiltration at the implant–tissue interface. These results suggest that sustained control of bacterial burden through mechanical renewal of the surface not only suppresses infection but also stabilizes host tissue responses, highlighting the importance of dynamic biointerface regulation in preventing chronic implant-associated infection.

Importantly, the actuation capability of DARTS is not limited to wired operation. Beyond electrical stimulation, DARTS also supports remote, noninvasive control, as demonstrated through NIR laser-triggered actuation. Localized photothermal activation of the embedded actuator enables transdermal deformation of implanted DARTS without physical connections, while maintaining physiologically relevant temperatures in surrounding tissues. This capability expands the translational relevance of DARTS by enabling on-demand, external control after implantation, a feature that is particularly attractive for clinical scenarios requiring periodic intervention without surgical access.

As a proof-of-concept platform, DARTS also presents several limitations and opportunities for future development. Actuation strategies may require optimization for different implant geometries, implantation depths, and anatomical environments. While both electrical and remote actuation were demonstrated here, implant-specific control schemes will be necessary for clinical translation. In addition, longer-term implantation studies are needed to assess chronic host responses, mechanical durability, and long-term stability of dynamic actuation under physiological conditions. Finally, although DARTS relies on purely physical antibacterial mechanisms, the platform is inherently modular and could be augmented with additional functionalities, such as immunomodulatory or catalytic effects, to further enhance performance against highly resilient or polymicrobial infections.

In summary, DARTS establishes dynamically actuated surface topography as a generalizable materials strategy for sustained control of biointerfaces. By decoupling antibacterial function from chemical release and introducing mechanical renewability at the surface, DARTS suppresses bacterial attachment, disrupts biofilms, and enhances therapeutic susceptibility over time. This approach provides a foundation for next-generation infection-resistant implants and mechanically adaptive biomaterial surfaces designed to operate reliably in dynamic biological environments.

## METHODS

### Ethical statement

All animal procedures were performed in accordance with institutional and federal guidelines for the care and use of laboratory animals. The mouse subcutaneous infection experiments were approved by the University of Michigan Institutional Animal Care and Use Committee (IACUC protocol #PRO00012380). No human subjects were involved in this study.

### Materials

Gentamicin sulfate (GS), anhydrous ethanol, acetone, dichloromethane (DCM), (tridecafluoro-1,1,2,2-tetrahydrooctyl)trichlorosilane, lithium fluoride (LiF), hydrochloric acid (HCl), and sulfuric acid (H_2_SO_4_) were obtained from Sigma-Aldrich. Polydimethylsiloxane (PDMS) base and curing agent (Sylgard 184) were sourced from Fisher Scientific. Ecoflex (00-50) and Dragon Skin (10) were obtained from Smooth-On. Ti_3_AlC_2_ MAX phase precursor powders (300 mesh, ≥99% purity) were supplied by Laizhou Kai Ceramic Materials Co. Shrinkable clear polystyrene (PS) sheets were purchased from Grafix. Deionized (DI) water (18.2 MΩ, Milli-Q system) was used throughout. Shape memory alloy (SMA, Nitinol Micro-Springs) was bought from Kellogg’s research labs. All materials were used as received without any additional purification steps.

### Synthesis of Ti_3_C_2_T_x_ MXene nanosheets

Ti_3_C_2_T_x_ MXene nanosheets were synthesized through an in situ hydrofluoric acid-generating etching process. Briefly, Ti_3_AlC_2_ MAX phase powder (1.0 g) was reacted with a mixture of LiF (7.5 g) and 9 M HCl (40 mL) at 35 °C for 24 h under continuous stirring to selectively remove the Al layers. The resulting solid was collected by centrifugation and repeatedly washed with deionized water until the suspension reached a pH of approximately 6.0. The partially delaminated material was then dispersed in 100 mL of DI water, bath-sonicated for 1 h at 4 °C, and centrifuged at 3500 rpm for 30 min to sediment the unexfoliated fractions. The resulting supernatant comprised stable Ti_3_C_2_T_x_ MXene nanosheets, with a final suspension concentration of roughly 10 mg/mL.

### Layer-by-Layer (LbL) assembly of MXene multilayers

MXene/GS multilayer coatings were assembled on various substrate types, including PS films, and glass slides. GS and Ti_3_C_2_T_x_ MXene dispersions were diluted to 1.0 mg/mL in DI water, with pH values of 5.0 and 6.1, respectively. The high charge density promotes self-assembly processes (Supplementary Fig. 1), especially, LbL self-assembly as it provides stronger electrostatic interactions between GS and MXene. All substrates were cleaned by sequential sonication in ethanol and water (15 min each), dried with nitrogen, and activated by oxygen plasma treatment (Tergeo-Plus, PIE Scientific) for 10 min at 50 W.

LbL deposition was performed using an automated dip-coating system (KSV NIMA Dip Coater, Biolin Scientific). Plasma-treated substrates were initially immersed in a PDAC solution for 30 min to form a positively charged foundation layer, followed by two DI water rinses (2 min each).

Multilayers were then built by alternating immersion in the cationic solution for 5 min, rinsing twice (2 min each), and then soaking in the anionic solution for 5 min with the same rinse sequence.

Each complete cycle produced one (MXene/GS) bilayer, and the process was repeated to obtain films with the desired number of bilayers, denoted as (MXene/GS)_n_. The resulting multilayer-coated substrates were air-dried at room temperature and stored in sealed containers until use.

MXene-coated PS sheets were cut to the desired dimensions and thermally shrunk in an oven at 130 °C for 5 min to generate wrinkled surface features. The shrinkage strain of the PS substrate was calculated using *ε* = (*L*ᵢ - *L*_f_)/*L*ᵢ. A degassed PDMS mixture (prepolymer:curing agent = 15:1 w/w) was then spin-coated onto the wrinkled MXene films at 500 rpm for 1 min, producing a layer approximately 300 µm thick, followed by curing at 60 °C for 4 h to obtain PS/MXene LbL/PDMS composites. These composites were subsequently submerged in dichloromethane (DCM) for 2 h to dissolve the PS, and the remaining structure was rinsed with acetone and ethanol. After drying at room temperature, MXene multilayers supported on PDMS were collected for further use.

### Fabrication of SMA-embedded elastomeric actuators

SMA spring was cut to a length of approximately 5.3 mm and electrically connected to sheath-stripped wires at both ends. The spring was then elongated to about 150% of its original length and secured within a custom 3D-printed mold. Elastomer precursors (Ecoflex and Dragon Skin) were prepared at a 1:1 mixing ratio and poured into the mold to encapsulate the stretched SMA. After curing for 4 h, the SMA-elastomer composite was removed from the mold, thoroughly rinsed with methanol, ethanol, and deionized water, and fully dried. The actuator was finalized by trimming excess material and cleaning the wire ends. Although the actuator dimensions can be scaled as needed, the standard size used in this study was 10 mm (length) x 1.5 mm (height) x 4 mm (width). For in vivo applications, exposed wire segments were coated with elastomer or replaced with enamel-insulated wires to prevent electrical leakage.

### Integration of MXene multilayers with SMA actuators

After fabricating the SMA-embedded elastomeric actuators and preparing the MXene multilayers on PDMS, the MXene films were bonded to the actuator using a thin layer of PDMS (prepolymer: curing agent = 15:1, w/w). The assembled device was then cured at 37 °C overnight to ensure complete adhesion.

### Electrical and remote laser actuation

For electrical actuation, the device was connected to a programmable power system consisting of a DC power supply (Model 1403, Global Specialties), a control circuit board, and a microcontroller (Metro 328, Adafruit). The microcontroller generated user-defined electrical signals to regulate actuation parameters such as duty cycle and frequency.

For remote activation, a NIR laser source with an 854 nm wavelength (LD852-SE600, Thorlabs) equipped with an adjustable collimating lens (LTN330-A, Thorlabs) was used. The laser was driven at 600 mA to achieve maximal output power and focused at a working distance of 5 cm. Laser intensity at various distances was determined by dividing the measured optical power (PM160, Thorlabs) by the beam spot area, the latter visualized using a visible-infrared detector card (VRC2, Thorlabs).

### Materials characterization

Film thickness and surface topography were characterized using atomic force microscopy (AFM; Dimension Icon, Bruker) operated in tapping mode. Optical absorbance during LbL deposition was monitored with a UV-vis spectrophotometer (Agilent 8453). For thickness measurements, MXene multilayers deposited on glass were gently scored with a plastic razor blade, and the step height was recorded at three predefined positions. Surface morphology of the assembled films was examined by SEM (Tescan Mira3).

### Bacterial culture and biofilm assays on static topography

*S. aureus* (USA300), *S. epidermidis* (AmMS 242), *P. aeruginosa* (AMC [NRRL B-12]), and *E. coli* (UTI89) were used for antibacterial and biofilm studies. Bacterial strains were first streaked onto agar plates and incubated overnight, followed by growth in LB broth at 37 °C with shaking at 200 rpm. Agar and LB media specific to each strain were prepared according to the manufacturer’s guidelines. Overnight cultures were diluted to approximately 10^4^ CFU/mL and applied onto sample surfaces to initiate biofilm formation. Wrinkled or planar substrates (1 × 1 cm^2^) were placed at the bottom of 24-well plates and incubated for 24 h (*S. aureus* and *S. epidermidis*) or 48 h (*P. aeruginosa* and *E. coli*) at 37 °C.

To quantify bacterial attachment, biofilms were stained with a mixture of SYTO 9 and propidium iodide (PI). Prior to staining, samples were rinsed three times with PBS. Surfaces were then incubated with SYTO 9 (5 µM) and PI (20 µM) for 15 min at room temperature in the dark.

Fluorescence imaging was performed using an Axio Observer microscope (Carl Zeiss), and quantitative image analysis was conducted with ImageJ and CellProfiler to distinguish live versus dead bacteria.

### SEM preparation of biofilms on MXene surfaces

Biofilms grown on MXene surfaces with static wrinkles were prepared as described above. Samples were rinsed with PBS and fixed in 4 % paraformaldehyde for 15 min, followed by an additional PBS wash. The specimens were then post-fixed in 1 % osmium tetroxide in PBS for 30 min. After fixation, samples were dehydrated through a graded ethanol series (30%, 50%, 70%, 100%, 100%), spending 5 min in each solution. Dehydrated samples were dried using a critical point dryer (Leica EM CPD300), sputter-coated with gold, and imaged using a scanning electron microscope (Tescan MIRA3).

### Kirby–Bauer disk diffusion assay

*S. aureus* was cultured on agar plates, and MXene-coated samples were placed without direct contact. Gentamicin disks served as positive controls. After incubation at 37 °C for 18-24 h, inhibition zones were evaluated.

### On-demand biofilm detachment via dynamic topography

To evaluate biofilm detachment induced by dynamic topography, biofilms were first grown on wrinkled MXene surfaces at 37 °C using an initial inoculation density of ∼10^4^ CFU/mL. Prior to actuation, all samples were gently rinsed with PBS. Actuators bearing the biofilms were then subjected to uniaxial stretching for 20 cycles over 6 min 20 s (0.0526 Hz) while submerged in PBS. Following dynamic actuation, samples were washed again with PBS, and the remaining adherent bacteria were stained and imaged using the same procedures described above. Biofilm coverage on actuated versus non-actuated surfaces was quantified using identical image analysis methods.

### Antibiotic susceptibility test

Benchmarking of antibacterial performance was performed using a combination of microdilution MIC testing, surface inoculation, dynamic actuation, antibiotic challenge, and CFU quantification. MICs were determined by standard broth microdilution in cation-adjusted Mueller-Hinton broth (CAMHB). Vancomycin was prepared as a two-fold serial dilution across columns 1-10 of a 96-well plate, beginning with 64 µg/mL (2x) and diluted to 0.5 µg/mL (final concentrations after inoculation: 32-0.25 µg/mL). Growth controls contained CAMHB with inoculum but no antibiotic, and sterility controls contained CAMHB alone. The bacterial inoculum was adjusted to ∼1.5 × 10^8^ CFU/mL in CAMHB (OD_600_ = 0.1). For each well, 100 µL of the 2x vancomycin dilution was combined with 100 µL of bacterial inoculum (or CAMHB for controls), yielding a total volume of 200 µL. Plates were incubated at 37 °C for 16-20 h, and the MIC was defined as the lowest concentration with no visible growth relative to the growth control.

Biomaterial surfaces representing four groups (Flat Static, DARTS Static, Flat Dynamic, and DARTS Dynamic) were inoculated with *S. aureus* suspensions diluted to ∼10^4^ CFU/mL and incubated for 24 h under appropriate growth conditions to allow surface colonization. Prior to actuation, each sample was gently rinsed with PBS. Dynamic samples were then placed in 1 mL PBS and subjected to uniaxial cyclic stretching for 20 cycles over 6 min 20 s (0.0526 Hz). After dynamic actuation, bacteria were collected from (i) the biomaterial surface (on surface) and (ii) the surrounding culture medium (in suspension) for subsequent CFU analysis.

Following actuation, samples were exposed to vancomycin at the concentration determined from the MIC assay and incubated for 16-20 h at 37 °C. After antibiotic treatment, surfaces were washed with PBS to remove residual drug. Bacteria remaining on the materials were recovered by vortexing and sonication. The supernatant containing the bacterial suspension was then collected. This suspension was serially diluted and plated onto agar, followed by ∼12 h incubation. CFUs were counted to quantify bacterial survival and removal efficiency for each experimental condition.

### In vivo implantation

The subcutaneous tissue of C57BL/6 mice was accessed by making a small skin incision, followed by gentle blunt dissection to separate the skin from the underlying muscle and create a pocket for biomaterial implantation. Two small exit sites were created for the electrical leads: one on the side opposite the incision and a second near the incision margin to accommodate the paired wires. The biomaterial construct was positioned directly on top of the muscle, and the overlying skin was gently pressed to ensure proper contact. The main incision was then closed using 6-0 Vicryl sutures. Digital photos of the step-by-step procedure of the implantation are shown in **Supplementary Fig. 9**.

### Bacteria collection

After 1 or 7 days, the implanted biomaterials were retrieved, and adherent bacteria were recovered by vortexing and sonication. The resulting suspensions were serially diluted and plated on agar for CFU counting.

### Histopathological evaluation

Tissue biopsy samples for histopathological evaluation were processed by the University of Michigan Unit for Laboratory Animal Medicine (ULAM) In Vivo Animal Core (IVAC). Briefly, samples were fixed in 10 % neutral-buffered formalin, paraffin-embedded, sectioned, and stained with H&E, Twort’s Gram stain, and Masson’s trichrome. All sections were evaluated and semi-quantitatively scored by a board-certified veterinary pathologist who was blinded to the experimental groups, according to the scoring rubric outlined in **Supplementary Table 1**.

### Statistical analysis

Statistical analyses were performed using GraphPad Prism version 10 (GraphPad Software Inc.). For comparisons between two groups, two-sided nonparametric t-tests were used. For comparisons involving more than two groups, one-way ANOVA followed by Tukey’s multiple comparisons test was used, unless otherwise noted. p-value < 0.05 was considered statistically significant. Details on data presentation, sample sizes, and statistical tests used are provided in the figure legends.

## ACKNOWLEDGEMENTS

The authors acknowledge financial support from the NIH R35GM157070 and the NSF 2441718 awarded to J.M., as well as the V Foundation V Scholar Award (V2023-002), which supported J.M.’s effort in part. S.N. gratefully acknowledges support from the Rogel Cancer Center Discovery Award, the Korean-American Scientists and Engineers Association (KSEA) Young Investigator Grant, and the NIH 1R03AG093279-01A1.

## DATA AVAILABILITY

All data needed to evaluate the conclusions of this study are included in the main text and/or the Supplementary Materials.

## AUTHOR CONTRIBUTION

J.M. M.A. and S.N. conceived the study. M.A. and J.L. performed experiments. M.A., J.M., and S.N. analyzed data. M.A., J.M., and S.N. wrote the manuscript, which was edited by all authors.

## Supporting Information

**Supplementary Figure 1.**
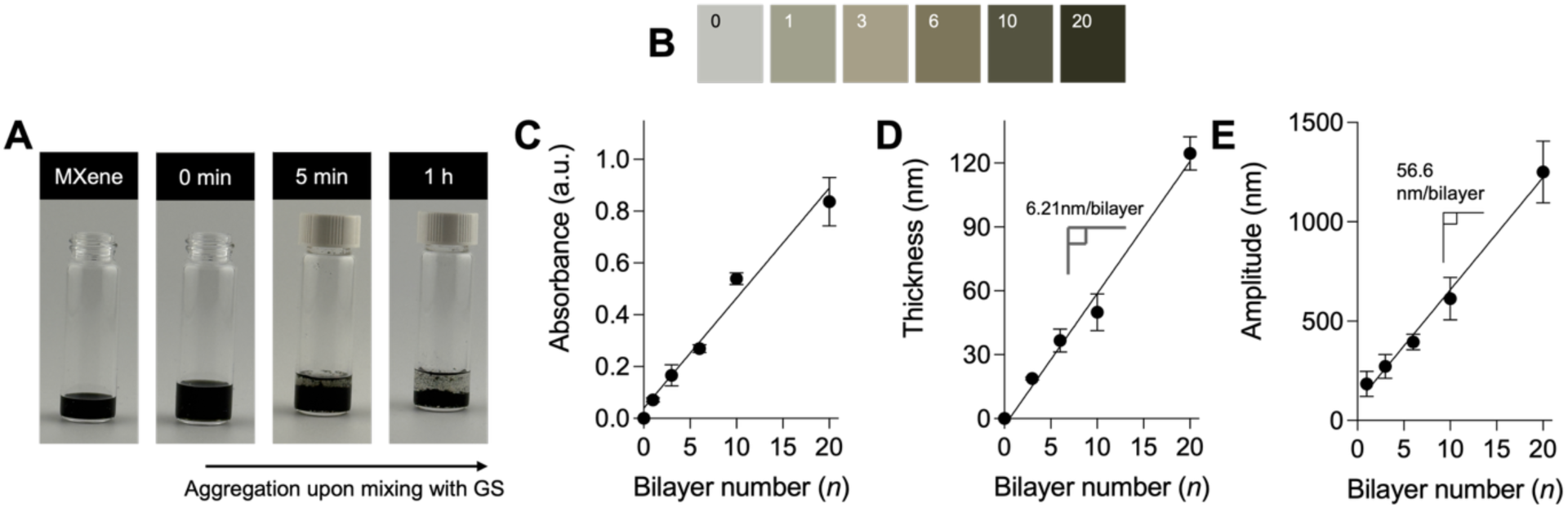
Characterization of MXene wrinkled topographies. **(A)** Optical images showing the time-dependent aggregation behavior of MXene dispersions upon mixing with gentamicin sulfate (GS). **(B)** Optical images of MXene multilayers with increasing bilayer number (n = 0, 1, 3, 6, 10, and 20). **(C)** UV–vis absorbance at 780 nm as a function of bilayer number (n = 5–9). **(D)** Film thickness measured by AFM as a function of bilayer number (n = 3–5). **(E)** AFM amplitude signal as a function of bilayer number (n = 3–4). Data are presented as mean ± S.D.

**Supplementary Figure 2.**
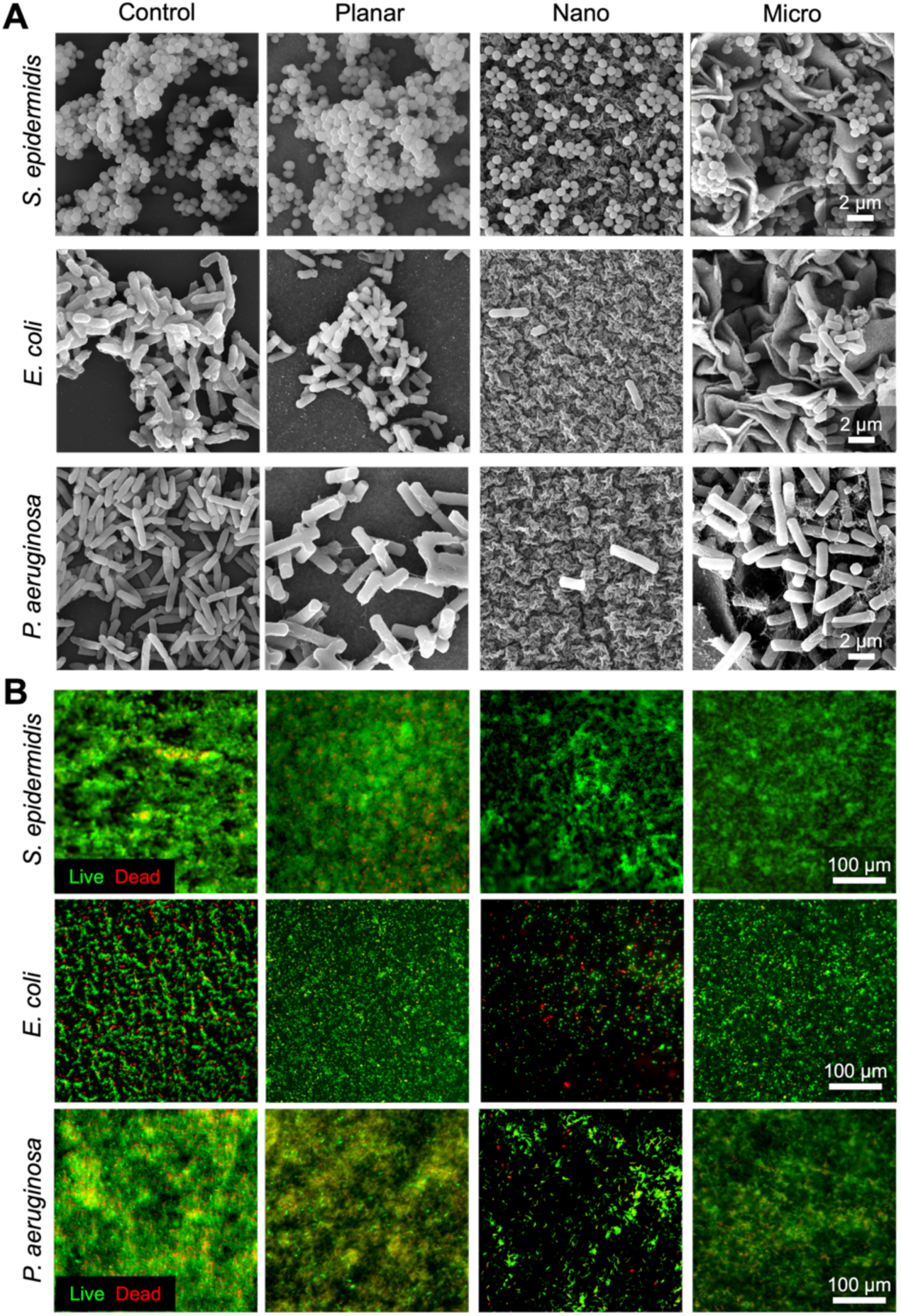
Antibacterial performance of wrinkled MXene topographies. **(A)** SEM and **(B)** fluorescence images of *S. epidermidis*, *E. coli*, and *P. aeruginosa* biofilms on Control, Planar, Nano, and Micro surfaces.

**Supplementary Figure 3.**
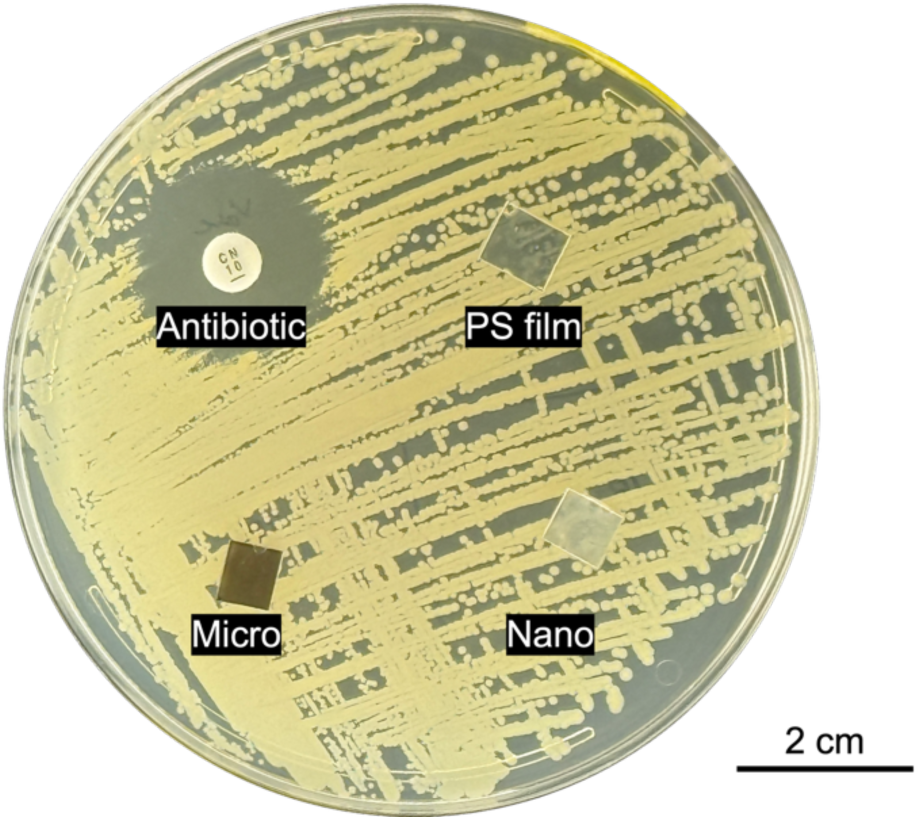
Kirby–Bauer disk diffusion assay. Representative agar plate showing bacterial growth in the presence of a gentamicin antibiotic disk, uncoated PS film, and MXene-coated Nano and Micro surfaces.

**Supplementary Figure 4.**
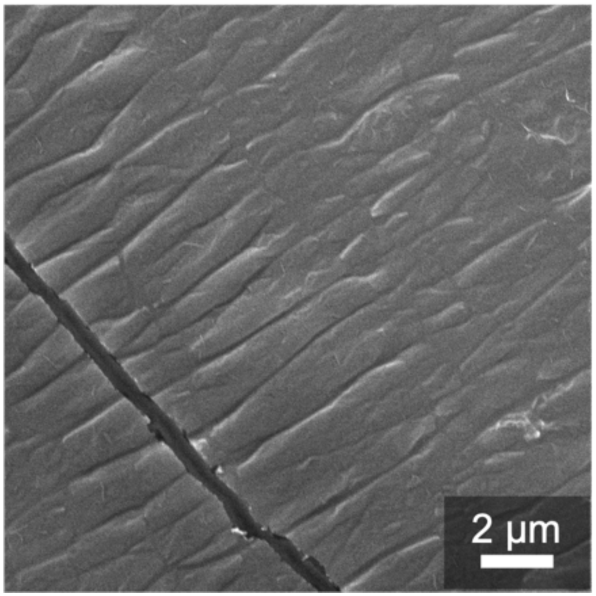
**A SEM image of MXene films directly assembled on PDMS.**

**Supplementary Figure 5.**
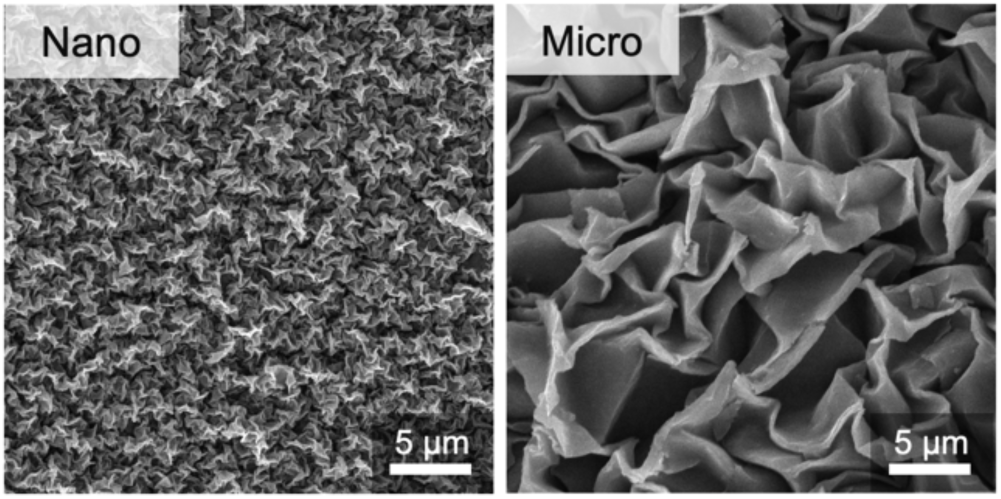
**SEM of nano- and micro-wrinkled MXene structures after transfer from PS films to PDMS.**

**Supplementary Figure 6.**
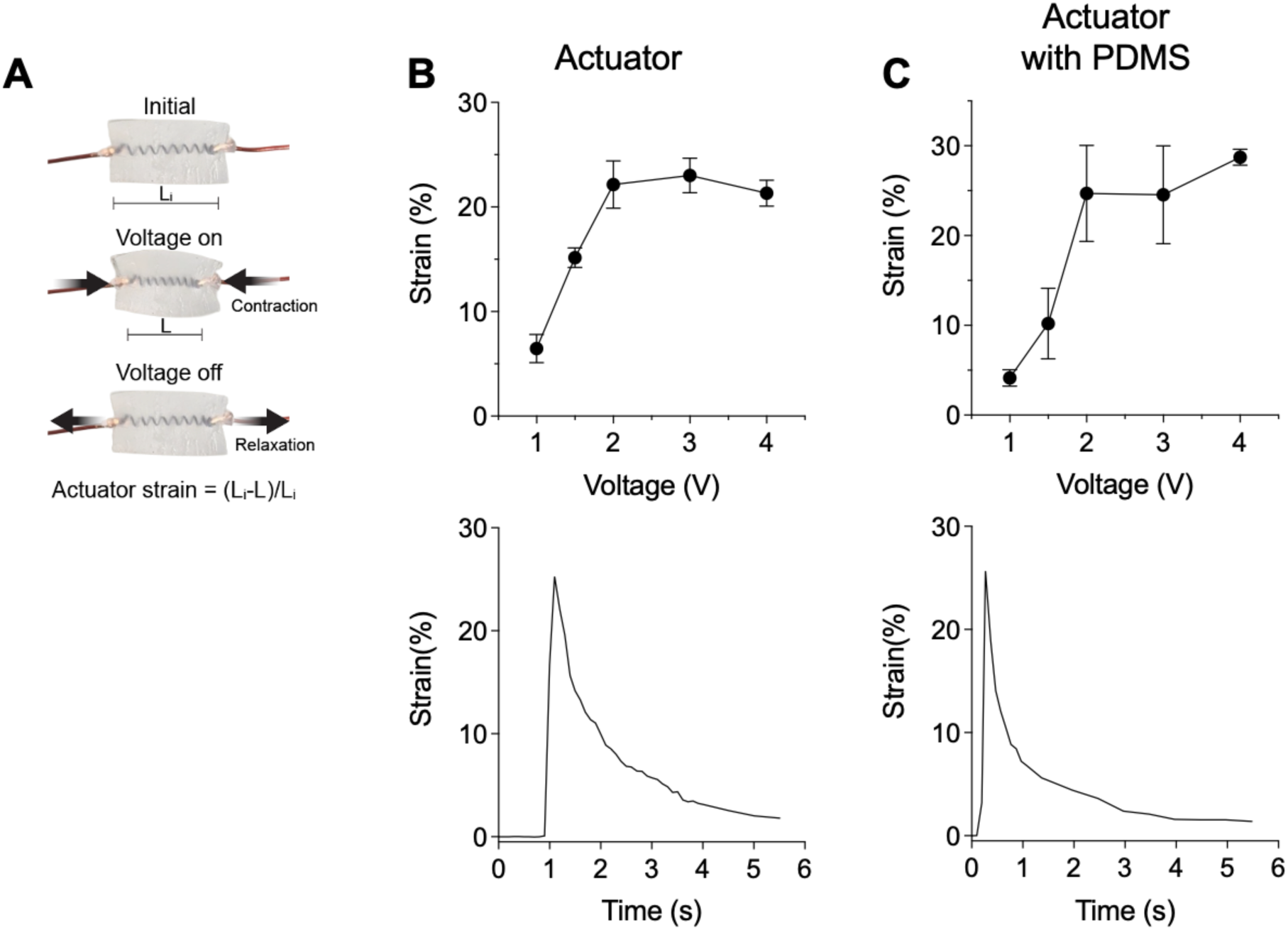
Mechanical actuation performance of the SMA actuator with and without PDMS support. **(A)** Schematic illustrating SMA-based actuation. **(B–C)** Actuation strain as a function of applied voltage and time (n = 5). Data are presented as mean ± S.D.

**Supplementary Figure 7.**
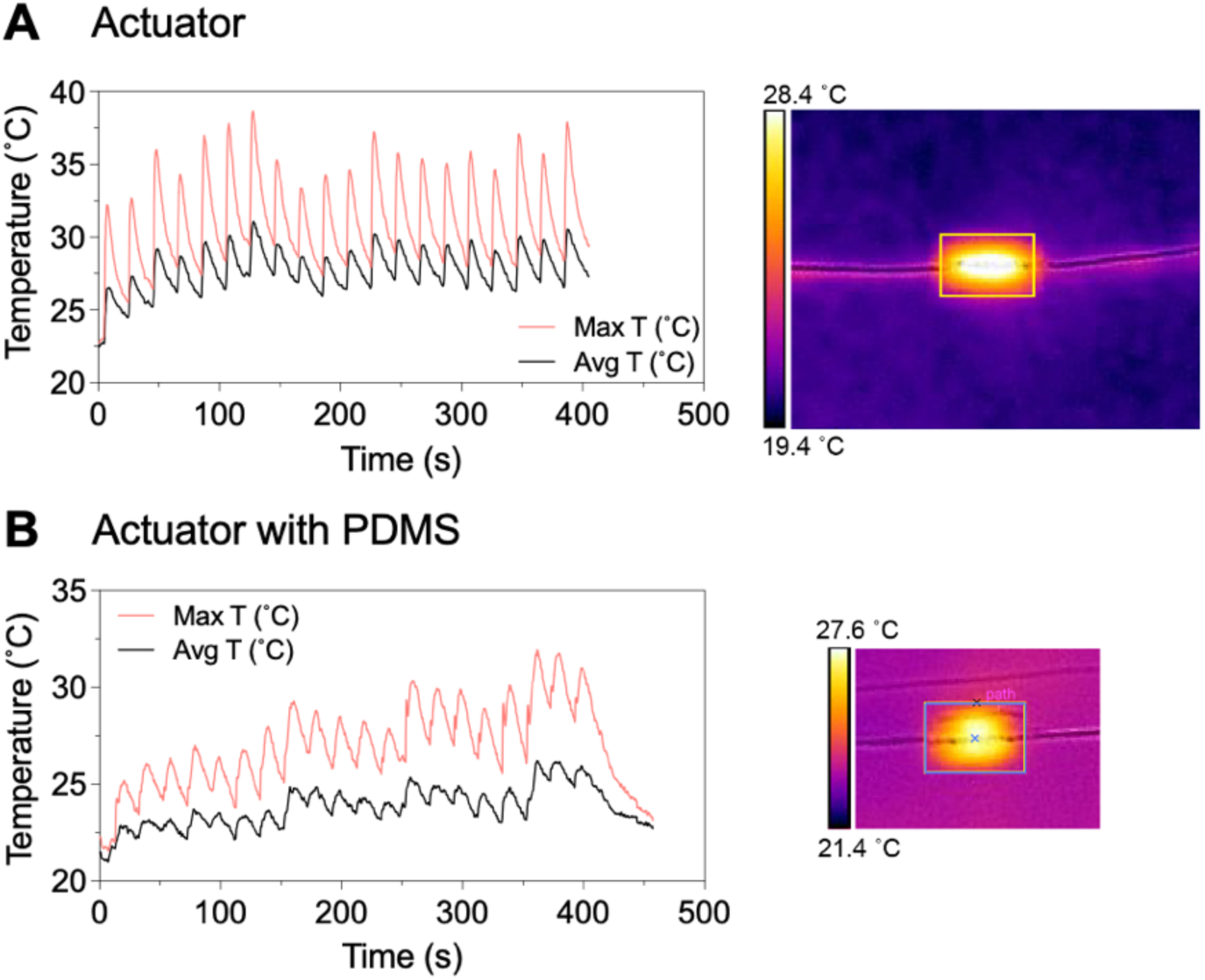
Thermal characterization of actuator. Actuation strain of actuators **(A)** with and **(B)** without PDMS as a function of applied voltage (n = 5), and infrared thermal imaging and corresponding temperature profiles during actuation.

**Supplementary Figure 8.**
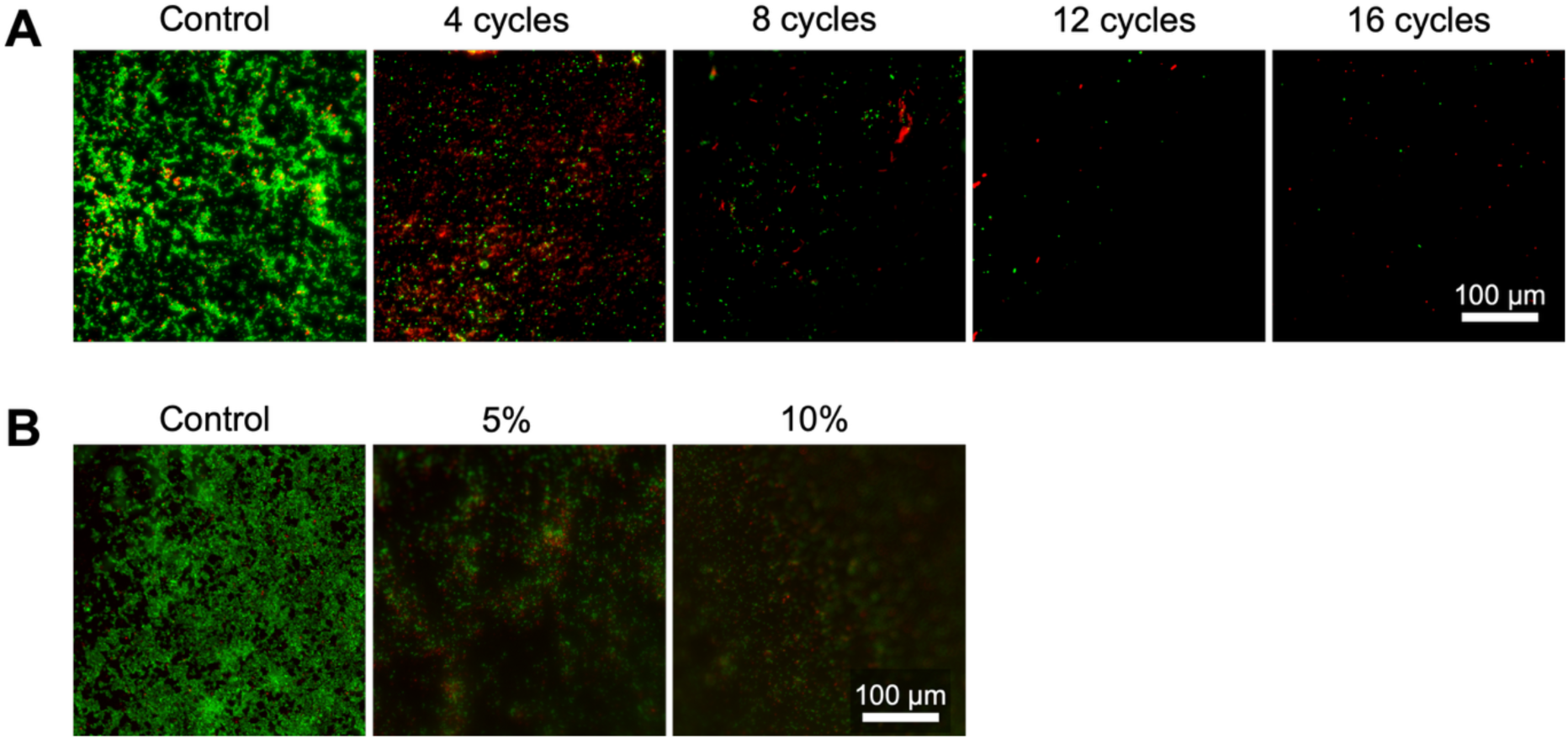
The effect of DARTS actuation parameters. **(A)** Representative live/dead fluorescence images of *S. aureus* biofilms subjected to increasing numbers of dynamic actuation cycles (0, 4, 8, 12, and 16 cycles), and **(B)** applied strain amplitude (0%, 5%, and 10%).

**Supplementary Figure 9.**
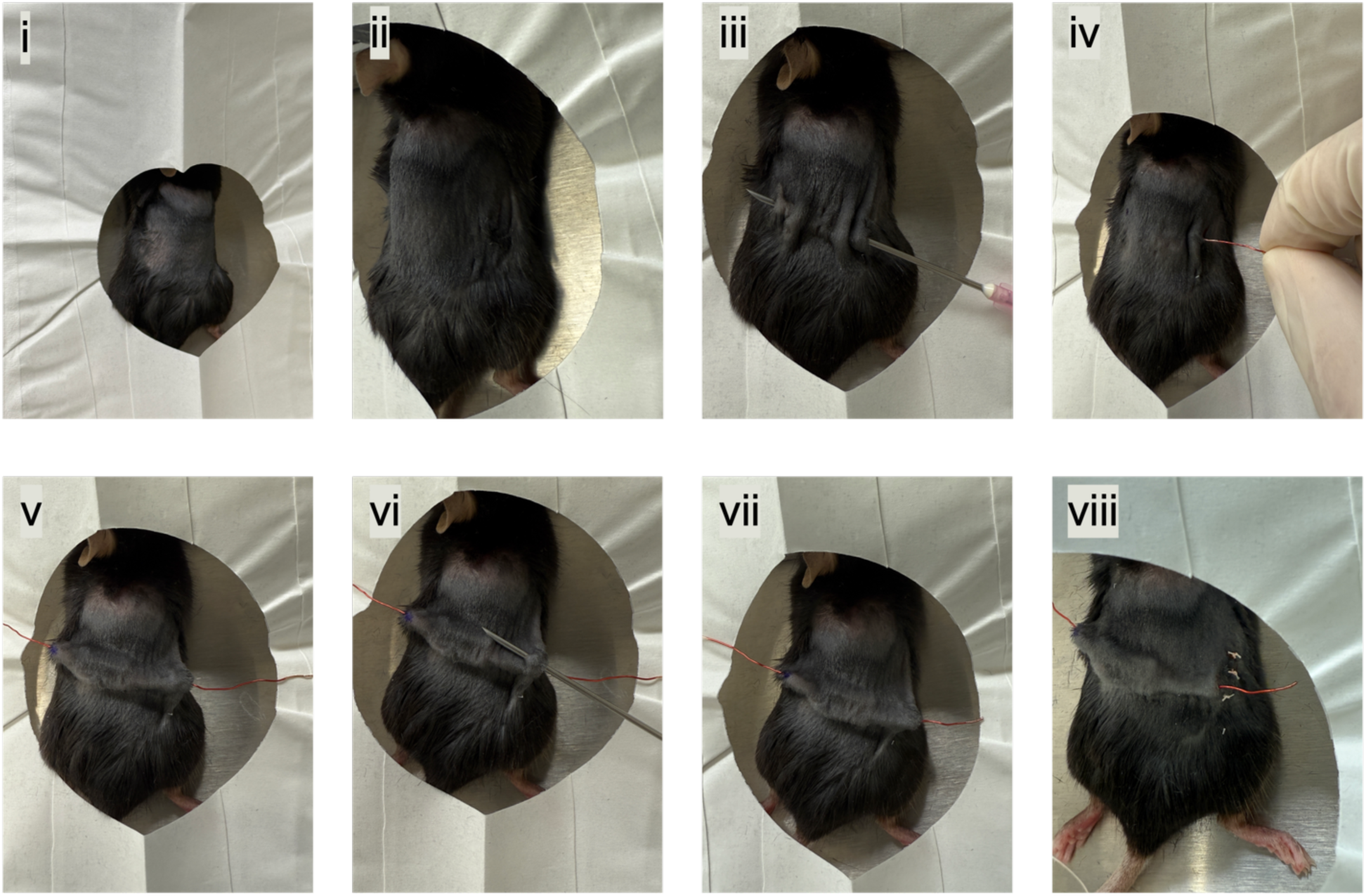
Digital photos of the surgery process. **(i)** Preparation of the surgical field following hair removal, sterilization, and positioning of the animal on a sterile surgical platform. **(ii)** Creation of a small dorsal incision using sterile surgical instruments. **(iii)** Formation of a small exit site on the left side using a needle for external wire placement. **(iv)** Introduction of the implant and associated wiring into the subcutaneous pocket. **(v)** Proper positioning of the implant beneath the skin with the left external lead aligned. **(vi)** Formation of a corresponding exit site on the right side using a needle for the right wire. **(vii)** Routing of the right external lead through the exit site. **(viii)** Closure of the incision site following implantation.

## Supplementary Note

Based on classical buckling theory^1–3^, the wrinkle wavelength (λ) of a stiff film on a compliant substrate is given by

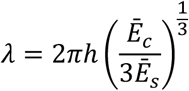

where ℎ is the film thickness, *E*_i_ is the elastic modulus, *v*_i_ is the Poisson’s ratio, and 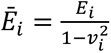 is the plane-strain modulus. For the coating layer composed of MXene multilayers, the Poisson’s ratio is *v*_c_ = 0.227,^4^ and the elastic modulus is *E*_c_ = 502 GPa,^5^ yielding a plane-strain modulus of *Ē*_c_ = 529.27 GPa. The substrate is a polystyrene (PS) film with *v*_s_ = 0.325,^6^ and *E*_s_ = 2 GPa,^6–9^ corresponding to *Ē*_s_ = 2.24 GPa. Substituting these values into the buckling equation gives *λ* = 26.95ℎ. For a single MXene layer (*n* = 1) with a thickness of ℎ = 9 nm, the predicted wavelength is *λ*_1_ = 242.55 nm, which is in good agreement with the experimentally measured value of 228 nm. Assuming linear scaling of wavelength with thickness, the ratio *λ*_1_/*λ*_20_ = ℎ_1_/ℎ_20_ = 9/125 predicts a wavelength of *λ*_20_ = 3000 nm for a 20-layer film, which closely matches the experimental value of 2700 nm. Although this buckling model is derived for a uniaxially pre-stretched substrate, it provides a reliable order-of-magnitude estimate and captures the correct thickness scaling for the biaxially stretched system used here.

---

**Supplementary Table 1.** Criteria and characteristics used for the histopathological analysis.

| **Epithelial damage** |  |
| --- | --- |
| 0 | Within normal limits |
| 1 | Mild, focal, superficial damage, serocellular crusting, with intact architecture and basement membrane |
| 2 | Multifocal to coalescing epidermal necrosis (ulceration) with exposure of disrupted basement membrane |
| 3 | Diffuse, full thickness of epidermal necrosis (ulceration) with obliteration of basement membrane |
| **Necrosis and tissue destruction** |  |
| 0 | Within normal limits |
| 1 | Mild, multifocal, necrosis in subcutis; dermis is intact with preservation of hair follicles and accessory glands |
| 2 | Severe necrosis predominantly in subcutis with minimal extension to dermis, and no to minimal damage on hair follicles and accessory glands |
| 3 | Marked coalescing necrosis in both subcutis and dermis, with obliteration and destruction of hair follicles and accessory glands; viable skin remains |
| 4 | Transmural diffuse necrosis manifested as non-viable skin at the implantation site |
| **Inflammation** |  |
| 0 | Within normal limits |
| 1 | Mild neutrophilic inflammation in subcutis |
| 2 | Severe neutrophilic inflammation in subcutis, with focal small extension to dermis |
| 3 | Severe, full thickness of inflammation composed of degenerate neutrophils, abundant cell debris, and large bacterial colonies |
| **Epidermal hyperplasia** |  |
| 0 | Within normal limits |
| 1 | Mild, simple hyperplasia |
| 2 | Moderate to marked, atypical hyperplasia to dyskeratosis |
| **Granulation** |  |
| 0 | Within normal limits |
| 1 | < 30% affected area |
| 2 | 30-70% affected area |
| 3 | >70% area |
| **Bacteria load** |  |
| 0 | No bacteria |
| 1 | Multiple small clusters of bacteria |
| 2 | Large volume of bacterial colonies |

